# Long-Read Structural and Epigenetic Profiling of a Kidney Tumor-Matched Sample with Nanopore Sequencing and Optical Genome Mapping

**DOI:** 10.1101/2024.03.31.587463

**Authors:** Sapir Margalit, Zuzana Tulpová, Tahir Detinis Zur, Yael Michaeli, Jasline Deek, Gil Nifker, Rita Haldar, Yehudit Gnatek, Dorit Omer, Benjamin Dekel, Hagit Baris Feldman, Assaf Grunwald, Yuval Ebenstein

## Abstract

Carcinogenesis often involves significant alterations in the cancer genome architecture, marked by large structural and copy number variations (SVs and CNVs) that are difficult to capture with short-read sequencing. Traditionally, cytogenetic techniques are applied to detect such aberrations, but they are limited in resolution and do not cover features smaller than several hundred kilobases. Optical genome mapping and nanopore sequencing are attractive technologies that bridge this resolution gap and offer enhanced performance for cytogenetic applications. These methods profile native, individual DNA molecules, thus capturing epigenetic information. We applied both techniques to characterize a clear cell renal cell carcinoma (ccRCC) tumor’s structural and copy number landscape, highlighting the relative strengths of each method in the context of variant size and average read length. Additionally, we assessed their utility for methylome and hydroxymethylome profiling, emphasizing differences in epigenetic analysis applicability.

## Introduction

One of the most prominent signs of carcinogenesis is the structural deviation of the cancer genome from that of the parent cell, which often involves large structural and copy number variations (SVs and CNVs, respectively). Such variations are annotated as structural features (deletions, insertions, duplications, inversions, and translocations) differing from the human genome reference or a matched sample for comparison ^1^. Traditionally, cytogenetic techniques such as karyotyping are used to detect large structural variations ranging from whole chromosome duplications, chromosome arm deletions, and down to SVs of ∼ 5-10 mega basepairs (Mb) ^2,3^. Applying fluorescence in-situ hybridization (FISH) techniques may bring the resolution down to several hundred kilo basepairs (kbp) ^4^, but a critical resolution gap remains between cytogenetics and short-read sequencing. The lack of access to genomic variation on the scales of 1-500 kbp has nourished the development of long-read technologies that can address this need. Various long-read methods have been introduced in recent years, including SMRT sequencing commercialized by PacBio, which routinely provides high-quality reads on the 10 kbp scale ^5^. Another concept involves a library preparation technique that allows linking proximal DNA fragments computationally by sequence barcode ligation (linked reads-10x genomics ^6^/TELseq ^7^). Here, we utilized two techniques, Optical genome mapping and ONT sequencing, that stand out in their ability to cover the full gap in mapping ability between short-read sequencing and karyotyping, offering an enhanced alternative to traditional cytogenetic analysis.

Optical genome mapping (OGM), commercialized by Bionano Genomics Inc. (BNG), has already gained clinical utility and is emerging as an alternative to cytogenetics by mapping the coarse grain structure of unamplified genomic fragments hundreds of kbp in length ^8,9^. The molecules are labeled at a specific sequence motif (CTTAAG) by a methyltransferase enzyme that transfers a fluorescent molecule to the labeling site from a synthetic cofactor analog. Every molecule acquires a sequence-specific fluorescent pattern along the DNA backbone during this process. The labeled DNA sample is applied to a silicon chip, where the molecules are electrophoretically extended in an array of parallel nanochannels. Millions of long, extended DNA molecules with their overlaying fluorescent barcode are imaged in the channels at high throughput. Once the images are digitized, DNA molecules may be mapped to their genomic location according to the pattern of fluorescent spots along the DNA and its matching to the expected pattern on the genome reference. Alternatively, the patterns may be stitched and assembled to build the whole genome structure *de-novo* ^10^.

Oxford Nanopore Technologies (ONT) is another prominent player in the long-read mapping and sequencing space. In recent years, ONT sequencing reads have gotten longer, error rates have diminished, throughput has increased, and prices per genome have dropped to levels that may justify clinical utility ^11,12^. For sequencing, DNA molecules are translocated through protein pores while measuring the electric ionic current flowing through the pore. Different sequence compositions generate various degrees of current attenuation, which is then computationally interpreted to generate the base sequence of the translocated DNA molecules. While offering single-base resolution, ONT provides shorter median read lengths compared to OGM. Both methods may be applied to native DNA that still carries chemical DNA modifications such as DNA methylation or DNA damage adducts. This gives rise to another beneficial feature: the acquisition of epigenetic information during genetic analysis. In OGM, an additional color may be used to chemically tag modifications of interest and create a hybrid genetic/epigenetic physical map of the molecules ^10,13–16^. ONT, on the other hand, does not require any additional preparative steps for calling epigenetic modifications as it relies solely on the electrical contrast generated by the native chemical structure of the modified base ^17^. Nevertheless, accurate modification calling requires a complete training set, which is not trivial for most base modifications.

Herein, we used both methods in order to characterize the structural and copy number landscape of a matched clear cell renal cell carcinoma (ccRCC) tumor-normal sample pair, pinpointing the specific strengths of each technique. Additionally, we characterized the methylome and hydroxymethylome of the pair and highlighted differences in practical utility for epigenetic analysis.

## Methods

### Patient clinically relevant information

Tumor and normal adjacent tissue were obtained in the course of radical nephrectomy performed in an 82-year-old male. Tumor was diagnosed histologically as ccRCC with morphological features of eosinophilic variant at pT3a stage. Tissues were stored from the time of surgery to analysis at -80°C (fresh-frozen sample).

Sample collection and handling was approved by institutional review boards in accordance with the declaration of Helsinki.

### Extraction of high molecular weight DNA

Ultra-high molecular weight (UHMW) DNA for 5-hmC OGM and ONT analyses was extracted using *SP Tissue and Tumor DNA Isolation kit* (Bionano Genomics), according to the manufacturer’s protocol. High molecular weight (HMW) DNA for OGM unmodified CpG analysis was extracted using *Animal Tissue Isolation kit* (Bionano Genomics) according to *Bionano Prep Animal Tissue DNA Isolation Soft Tissue/Fibrous Tissue Protocol* for normal tissue/tumor, respectively.

### Nanopore sequencing (Oxford Nanopore Technologies)

Samples were prepared for sequencing using Ligation Sequencing Kit V14 (SQK-LSK114, Oxford Nanopore Technologies, UK) according to protocol with a starting DNA amount of 1 µg. Whole genome sequencing was performed on a “P2-Solo” device using R10.4.1 Flow cells (FLO-PRO11, Oxford Nanopore Technologies).

Basecalling of raw POD5 files was performed using the ONT proprietary software Dorado (v 0.3.2, Oxford Nanopore Technologies; https://github.com/nanoporetech/dorado) with the model: “dna_r10.4.1_e8.2_400bps_hac_@”. Reads were then aligned to the hg38 human reference genome using minimap2 ^18^ (v.2.24). Bam output files were then merged, sorted and indexed using samtools ^19^ (v1.16.1). SVs, CNVs and methylation and hydroxymethylation locations were called by the “wf-human-variation” pipeline (https://github.com/epi2me-labs/wf-human-variation) via EPI2ME software ^20^ (Oxford Nanopore Technologies) with minimum bam coverage set to 5. The default behavior of the pipeline is to report methylation and hydroxymethylation per CpG positions and with combined strands. Analyses were performed on a Linux operating system (Ubuntu 22.04.3) with Nvidia’s RTX 6000 GPU.

### Optical Genome Mapping (OGM)

#### a. Labeling DNA for genetic and 5hmC analysis

To create the genetic barcode, 750 ng of UHMW DNA in two reaction tubes were each mixed with 5X DLE-buffer (to a final concentration of 1X), 1.5 µL of 20X DL-Green and 1.5 µL of DLE-1 enzyme (Bionano Genomics) in a total reaction volume of 30 µL. The reaction was incubated for 4 hours at 37°C. Then, 5hmC sites were labeled by the enzyme β-glucosyltransferase from T4 phage (T4-BGT) ^15^. Magnesium chloride was added to 30 µL of DLE-labeled DNA to a final concentration of 9 mM. Then, the DNA was added to 4.5 µL of 10X NEBuffer 4 (New England Biolabs), uridine diphosphate-6-azideglucose (UDP-6-N3-Glu; (21)) in a final concentration of 50 µM, 30 units of T4 β-glucosyltransferase (New England Biolabs) and ultra-pure water in a final volume of 45 µL. The reaction mixture was incubated overnight at 37°C. The following day, dibenzocyclooctyl (DBCO)-ATTO643 ^21^ was added to a final concentration of 150 µM and the reaction was incubated again at 37°C overnight. The next day, the reaction tubes were added 5 µL of PureGene Proteinase K (Qiagen) and incubated for additional 30 minutes at 50°C. After the Proteinase K treatment, the two identical reaction tubes were merged and drop-dialyzed as one against 20 mL of 1X TE buffer (pH 8) with 0.1 µm dialysis membrane for a total of 2 hours. Finally, 300 ng recovered dual-color DNA was stained to visualize DNA backbone, by mixing it with 4X Flow Buffer (Bionano Genomics) to a final concentration of 1X, 1M DTT (Bionano Genomics) to a final concentration of 0.1 M, Tris (pH 8) to a concentration of 25 mM, NaCl, to a concentration of 25 mM, EDTA to a final concentration of 0.008-0.01 M, DNA Stain (Bionano Genomics) to a final vol/vol ratio of 8%, and ultrapure water. The reaction mixture was shaken horizontally on a HulaMixer for an hour and then incubated overnight at 4°C.

#### b. Labeling DNA for unmethylation analysis (reduced representation of unmodified cytosines in CpG context)

To create the genetic barcode, 1 µg of U/HMW DNA was mixed with 5X DLE-buffer (to a final concentration of 1X), 2 µL of 20X DL-Green and 2 µL of DLE-1 enzyme (Bionano Genomics) in a total reaction volume of 30 µL for 4 hours at 37°C, immediately followed by heat inactivation at 80°C for 20 minutes. Heat inactivation at these conditions degrades over 97% of the DL-Green cofactor, therefore preventing it from being incorporated by M.TaqI in the following reaction, and making the two reactions orthogonal. Then, unmodified cytosines in the recognition sequence TCGA were fluorescently labeled to perform reduced representation optical methylation mapping (ROM) ^14,22^. Two 500 ng reaction tubes of DLE1-labeled DNA were each mixed with 4 µL of 10X CutSmart buffer (New England Biolabs), 60 µM of lab-made synthetic AdoYnATTO643 ^21^, 1 µL of M.TaqI (10 units/µL; New England Biolabs) and ultrapure water in a total volume of 40 µL, and incubated for 5 hours at 65°C. Then, 5 µL of Puregene Proteinase K (Qiagen) were added and the reaction tube was incubated for additional 2 hours at 45°C. After the Proteinase K treatment, the two 500 ng reaction tubes were merged and drop-dialyzed as one against 20 mL of 1X TE buffer (pH 8) with 0.1 µm dialysis membrane for a total of 2 hours. Finally, 300 ng recovered dual-color DNA were stained to visualize DNA backbone by mixing it with 15 µL of 4X Flow Buffer (Bionano Genomics), 6 µL of 1M DTT (Bionano Genomics), 3 µL of 0.5M Tris (pH 8), 3 µL of 0.5M NaCl, 4.8 µL of DNA Stain (Bionano Genomics) and ultrapure water to a total volume of 60 µL, and incubated overnight at 4°C.

#### c. Running OGM

Labeled samples were loaded on Saphyr chips (G1.2) and run on a Saphyr instrument (Bionano Genomics) to generate single molecule maps. Optical mapping data from several runs were merged to a single dataset using Bionano Access (v1.6.1) and Bionano Solve (v3.6.1) (Bionano Genomics). The assigned channels for genetic and epigenetic labels in the molecules (.BNX) files were swapped with Bionano Solve (v3.6.1) according to manufacturer’s advice. *De novo* assemblies and “variant annotation pipeline” (single sample mode) for SV annotation were generated from 5hmC-labeled data with default parameters for human genomes using Bionano Access v1.7.1 and Bionano Solve v3.7.1. The *in-silico* digested human genome GRCh38 (*hg38_DLE1_0kb_0labels.cmap*) was used as the reference.

#### d. Epigenetic data processing

Molecules spanning over 150 kbp were aligned to the *in silico* human genome reference GRCh38, based on DLE-1 recognition sites (hg38_DLE1_0kb_0labels.cmap) using Bionano Access (v1.6.1) and Bionano Solve (v3.6.1), with default parameters according to the following combination of arguments: haplotype, human, DLE-1, Saphyr. Only molecules with an alignment confidence equal to or higher than 15 (P <= 10-15) that at least 60% of their length was aligned to the reference were used for downstream analysis. Alignment outputs were converted to global epigenetic profiles (bedgraph files) according to the pipeline described by Gabrieli et al. ^15^ and Sharim et al. ^14^ and in ebensteinLab/Irys-data-analysis on Github. Only regions covered by at least 20 molecules were considered.

### CNV analysis

In Order to generate CNV plots of OGM data, the coverage of DLE-1 labeling sites was extracted from raw output of CNV analysis (*cnv_rcmap_exp.txt*). Genomic regions with very high variance in coverage across Bionano Genomics’ control datasets compared to typical loci (hg38_cnv_masks.bed) were subtracted from analysis. Then, the mean coverage of such sites in 500,000 bp bins was calculated using Bedtools ^23^ *map* (v2.26.0). Then, for each bin, the log_2_ of the copy ratio (in a diploid organism, copy number/2) was calculated and plotted along the chromosomes. log_2_ of the copy ratio in 500,000 bp bins along ONT data was inferred by employing the EPI2ME workflow “wf-human-variation” to each sample. A running median over 10 bins was calculated to plot a smooth red line across the log_2_ of the copy ratio dots of both methods.

### SV analysis

Genomic coordinates, SV type and size of high-confidence (confidence score ≥ 0.5) annotated OGM SVs from “variant annotation Pipeline”, were extracted from output smap file and converted to bed format for downstream analysis. In case no end coordinate was supplied, it was taken as start+1. Both translocation breakpoints were considered for overlap with ONT SVs, but were counted as one event of a large SV (>10 kb). Only unique SVs were kept. SVs overlapping BNG’s list of N-base gaps in the reference or putative false positive translocation breakpoints (for “*de novo* assembly”, Solve 3.6.1) were masked from analysis (Bedtools *intersect -v* (v2.26.0)). Coordinates of OGM-detected SVs were extended by 500 bp up and downstream (Bedtools *slop* (v2.26.0) to account for possible differences in SV resolution between OGM and ONT. This extension was not considered to determine SV size.

Genomic coordinates, SV type and size of “passed” ONT SVs called by the EPI2ME workflow “wf-human-variation”, with minimum 5 reads supporting them, residing on canonical chromosomes, were extracted from the Sniffles2 VCF output file and converted to bed format for downstream analysis. End coordinate was taken as end+1 to avoid cases of 0 difference. Only unique SVs were kept. SVs overlapping BNG’s list of N-base gaps in the reference or putative false positive translocation breakpoints (for “*de novo* assembly”, Solve 3.6.1) were masked from analysis (Bedtools *intersect -v* (v2.26.0)). Only SVs of at least 500 bp were considered.

Overlap between SVs detected by OGM and ONT was calculated with Bedtools *intersect* (v2.26.0). The number of overlapping SVs is reported based on the ONT calls. Overlapping and non-overlapping SVs were then divided based on their size (absolute value).

Clinical significance analysis of ONT SVs was performed with SnpSift ^24^ against the ClinVar ^25^ database and had no findings.

### Alternative ONT SV callers

SV calling with SVIM ^26^ (https://github.com/eldariont/svim) was conducted with parameter: -- min_sv_size 50. The output VCF file was processed as described for the EPI2ME (Sniffles2) VCF. Only SVs of at least 500 bp were considered.

SV calling with CuteSV ^27^ (https://github.com/tjiangHIT/cuteSV) was conducted with parameters recommended for ONT data: --max_cluster_bias_INS 100; -- max_cluster_bias_DEL 100; --diff_ratio_merging_INS 0.3; --diff_ratio_merging_DEL 0.3; and additional parameters: -l 50; --min_support 5. The output VCF file was processed as described for the EPI2ME (Sniffles2) VCF. Only SVs of at least 500 bp were considered.

### Global epigenetic levels

Due to resolution differences between OGM and ONT, the mean epigenetic levels in non-overlapping 1000 bp genomic windows (generated using Bedtools *makewindow* (v2.26.0) was calculated. Only windows on canonical chromosomes that contain at least one relevant recognition site were considered (CpG for 5hmC and ONT mC, TCGA for OGM unmethylation; sites loci were extracted using the R package BSgenome (https://bioconductor.org/packages/release/bioc/html/BSgenome.html). To match the reported measure between OGM and ONT in methylation calling, the unmethylation level (1 – methylation level) was calculated from ONT methylation level. The weighted mean of all ONT epigenetic signals and ONT unmethylation signals in TCGA sites only (crossed with Bedtools *intersect* (v2.26.0)) in these genomic windows was calculated using Bedops *bedmap* ^28^ (v2.4.41). The number of OGM epigenetic labels and molecules covering each genomic window were counted using Bedtools *intersect* (v2.30.0). The average labels-to-molecules ratio across all windows was reported as the global epigenetic level for OGM.

To create bedgraphs of OGM signals, epigenetic labels and molecules were extended by 500 bp up and down stream to account for optical resolution ((Bedtools *slop* (v2.26.0)), prior to calculating the labels to molecules ratio in each genomic location. To reduce the resolution of OGM and ONT bedgraphs to 1 kbp windows, the weighted mean of signal in these windows was calculated with Bedops *bedmap* (v2.4.41).

### Gene expression data

Publicly available RNA-seq data of three tumor-matched pairs of ccRCC (stage 3) patients (PRJNA396588, GEO accessions: pair 1: GSM2723919, GSM2723920; pair 2: GSM2723927, GSM2723928; pair 3: GSM2723929, GSM2723930; ^29^) were aligned to the human genome (hg38) using TopHat ^30^ (v2.1.0) with default parameters and library-type and fr-firststrand flags, after retrieving the raw files with NCBI SRA toolkit ^31^. Only uniquely mapped reads were analyzed (minimal mapping quality of 30). Gene counts were obtained using HTSeq ^32^ (htseq-count, v0.11.3) against the GENCODE ^33^ (v34) reference gene models. Transcripts per million (TPM) scores were calculated.

### Epigenetic modifications signal along aggregated genes

Transcription start and end sites (TSS and TES) of protein-coding genes were defined according to GENCODE annotation (v34). Protein-coding genes were divided into four groups based on their average normalized TPM score in the RNA-seq of three matched ccRCC pairs. Unexpressed genes were defined as genes with TPM value <= 0.01 (∼3000 genes). The other expression groups are three equal quantiles of the expressed protein-coding genes (∼6000 genes per group). Mean 5-hmC and unmethylation signals along aggregated genes were calculated using DeepTools ^34^ *computeMatrix* (v3.5.4) in scale-regions mode, where each gene (from TSS to TES) was scaled to 15 kbp and divided into 300 bp bins. Compressed matrix output was summarized by DeepTools *plotProfile*. The average signal intensities for both markers were then plotted as a function of the scaled distance relative to the TSS.

## Results

We began by analyzing the genetic makeup of a stage 3 ccRCC tumor, a common type of kidney cancer known for characteristic structural abnormalities^35,36^, and a normal adjacent tissue. Our workflow consisted of extracting high molecular weight DNA, followed by per-protocol OGM and ONT analyses (**Fig. 1**).

**Figure 1.**
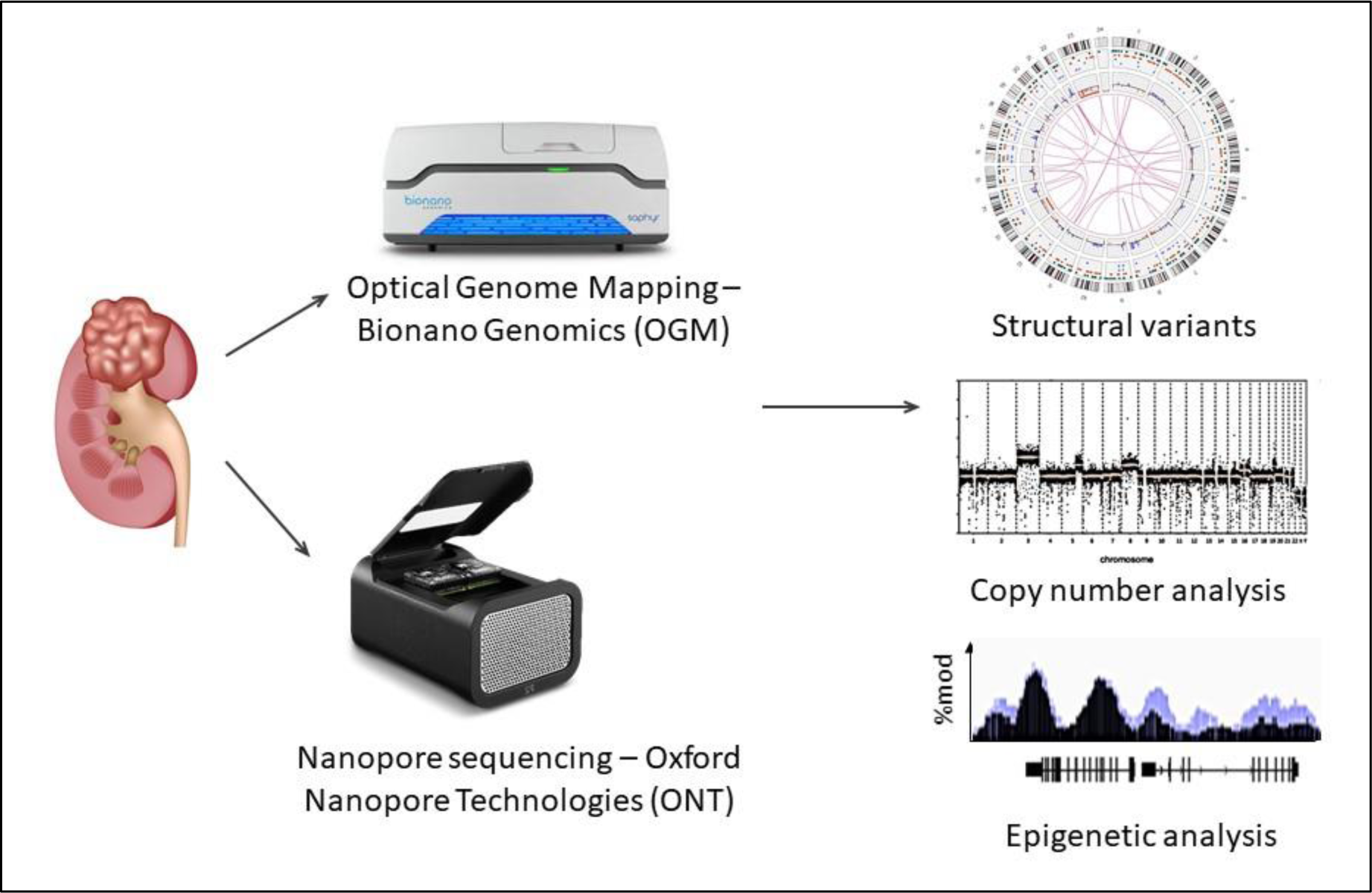
Experimental workflow. High molecular weight DNA was extracted from a ccRCC tumor and a normal adjacent kidney tissue. Samples were analyzed by OGM and ONT to detect structural and copy number variations, and epigenetic modifications. Results from both methods were compared.

By generating long reads, both methods unlock access to intricate areas of the genome, enabling the study of diverse structural variations, copy number variations and repetitive elements. Additionally, as both methods read native, unamplified DNA, they are able to detect epigenetic modifications. The different attributes of each technique affect their performance in the aforementioned analyses. **Table 1** is based on public company material and summarizes some of the performance specifications, pointing to advantages and limitations of the two methods and indicating their compatibility of use, depending on research goals and budget.

**Table 1.**
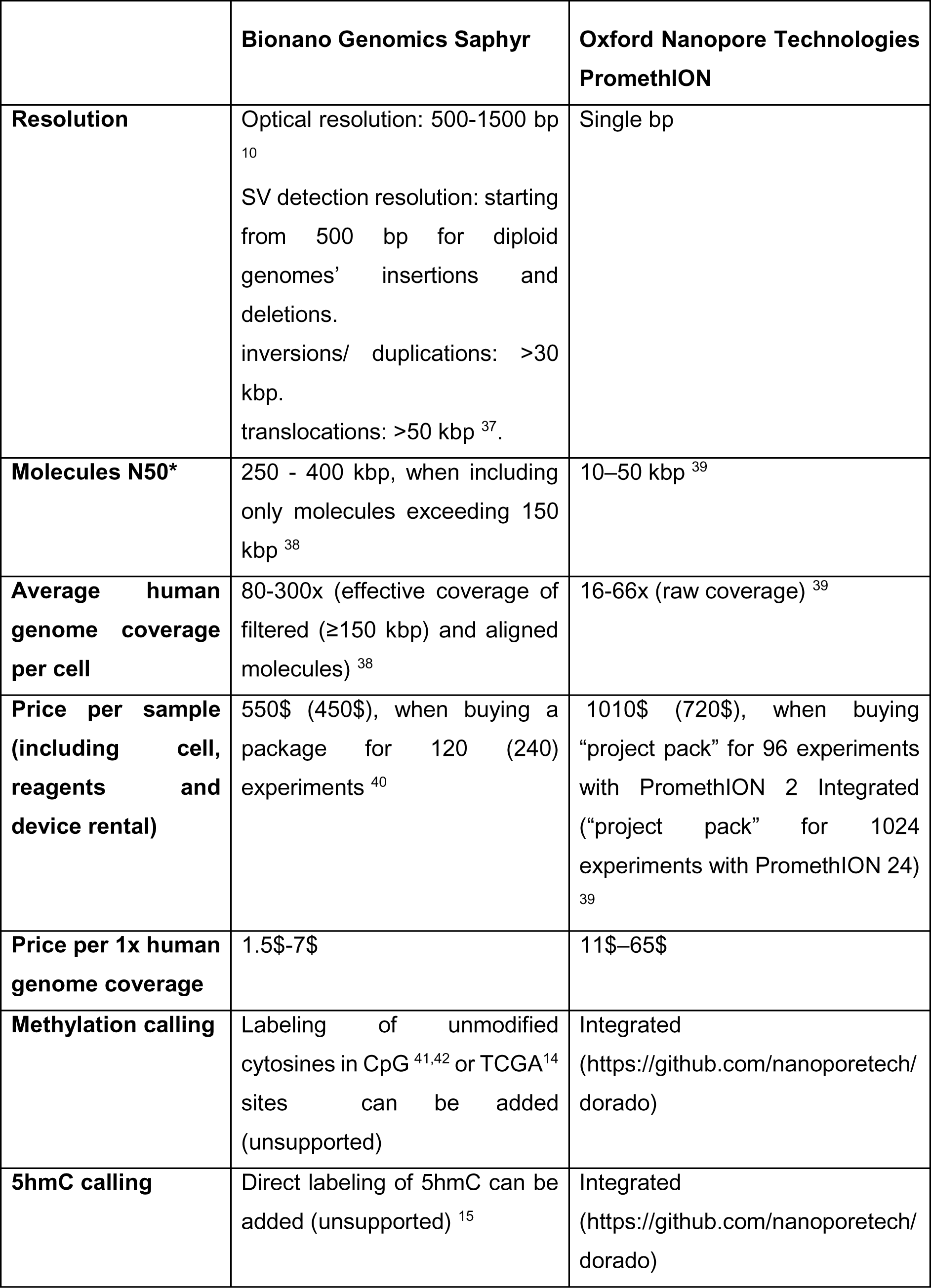
ONT and OGM specifications. * Molecules N50 is a measure of reads length indicating that half of the genetic data recorded came from reads longer or equal to this value.

### SV and CNV analysis of a ccRCC tumor

Clear cell renal cell carcinoma (ccRCC) is the most common type of renal carcinoma, and its incidence has been increasing in recent years. Over 90% of ccRCC cases demonstrate distinctive changes to the short arm of chromosome 3 (3p), from translocations and deletions to the loss of the entire chromosomal arm. Most cases involve the genetic or epigenetic inactivation of the von Hippel–Lindau (*VHL*) gene, located on this arm ^35,43,44^. Other frequently observed copy number variations and cytogenetic abnormalities in ccRCC include a gain of chromosome 5q, loss of 14q, trisomy of chromosome 7, loss of 8p, loss of 6q, loss of 9p, loss of 4p and loss of chromosome Y in men. Some CNVs were correlated with prognosis ^35,36,45^.

To compare the efficacy of the structural profiling and data analysis processes offered by each method, we applied CNV and SV analyses on data generated by both methods, adhering to the manufacturer’s recommended pipelines unless specified otherwise (see methods section). Herein, DNA from a ccRCC tumor and a normal adjacent tissue was analyzed using both ONT and BNG platforms. **Table 2** summarizes the resulting N50 and average coverage.

**Table 2.** Coverage and N50 of OGM and ONT genetic experiments.

|  | Effective genome coverage of aligned molecules | N50 of aligned molecules |
| --- | --- | --- |
| **OGM** | Tumor sample: 133X<br>Normal adjacent sample: 123X | Tumor sample: 291 kbp<br>Normal adjacent sample: 233 kbp |
| **ONT** | Tumor sample: 36X<br>Normal adjacent sample: 19X | Tumor sample: 18 kbp<br>Normal adjacent sample: 15 kbp |

First, genome-wide copy number, calculated in 500 kbp bins, was compared. Tumor plots are shown in **Figure 2.A** and normal adjacent tissue plots are shown in **Figure S1.** A running median over 10 bins was calculated to plot a smooth red line across the copy number dots. As expected, both methods produced highly similar CNV plots, identifying the loss of one copy of the entire 3p chromosomal arm, as well as a large DNA gain in 5q, and a smaller DNA loss in the same arm. Aneuploidies were found by both methods in chromosomes 7 and 12. OGM spotted a small DNA loss in chromosome 9 not reported by ONT. The normal adjacent sample did not exhibit any large copy number variation in both methods, suggesting somatic aberrations. The loss of 3p, gain in 5q and trisomy in chromosome 7 are well-documented genetic characteristics of ccRCC ^36,45^. The CNV plot generated with ONT data is smoother than with OGM due to resolution differences between the methods, influencing the number of data points sampled in each bin.

**Figure 2.**
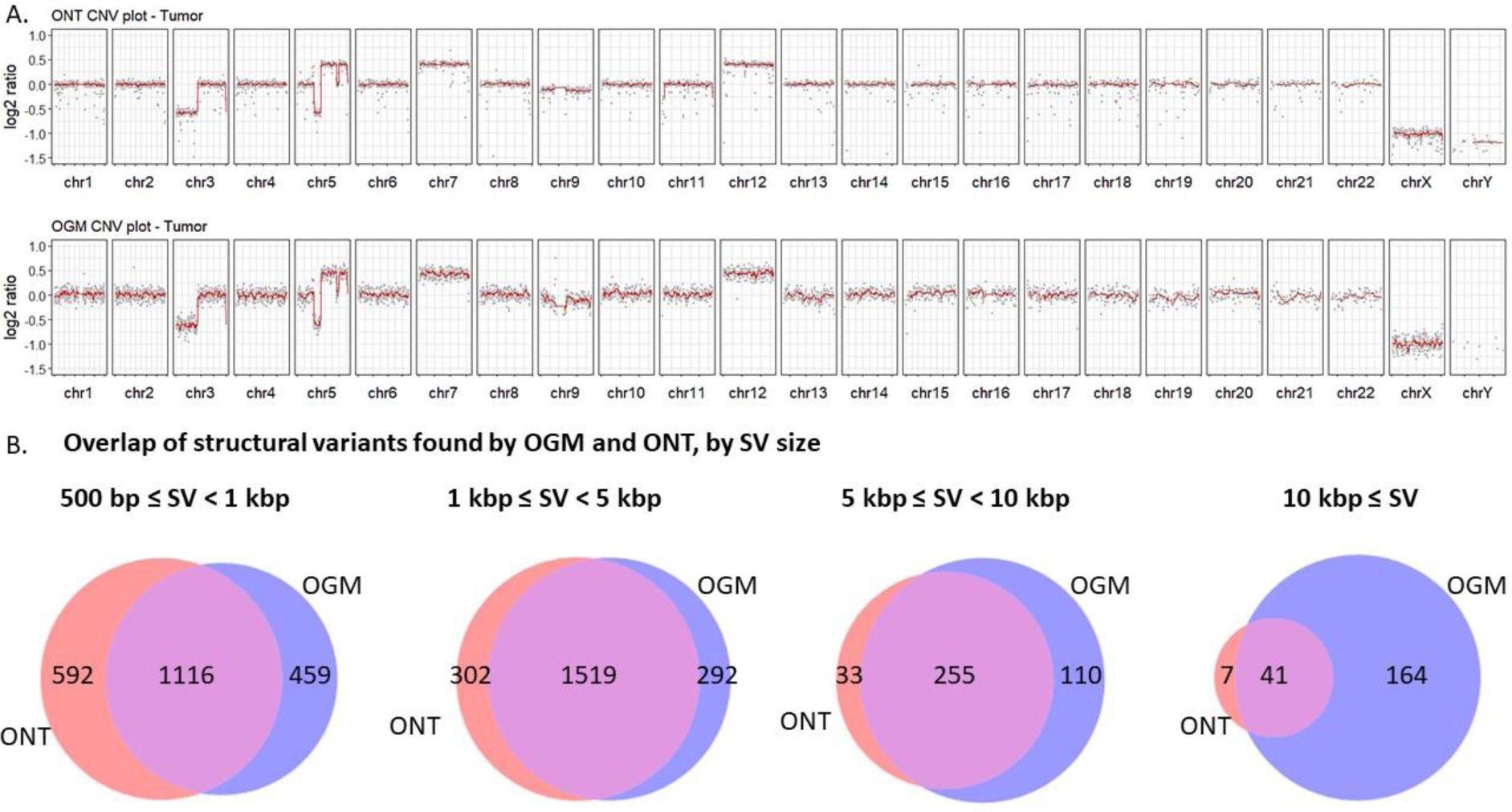
Comparative analysis of structural variations (SVs) and copy number variations (CNVs) in a ccRCC tumor, as detected by ONT and OGM. **A.** CNV plots (log_2_ of the copy ratio) generated from ONT (top) and OGM (bottom) data. Data illustrate highly similar findings, pinpointing a significant DNA loss on chromosome 3 and various losses and gains on chromosomes 5, 7 and 12. **B.** Venn diagrams displaying common and unique SVs to OGM and ONT, in four size ranges: 500 bp -1 kbp, 1-5 kbp, 5-10 kbp and above 10 kbp.

Clearly, both methods are adequate for basic Karyotyping; however, SV detection exhibited less congruence between the two methods. To facilitate a comprehensive comparison, we categorized detected SVs larger than 500 bp based on their size (**Figure 2.B**). As anticipated, for short SVs (500 bp - 1 kbp), ONT detected more SVs, leveraging its superior resolution at the base-pair level. In the intermediate SV length range (1 to 5 kbp), both methods demonstrated a comparable number of detections, with substantial overlap. Notably, OGM exhibited an advantage in the detection of larger SVs (more than 5 kb) owing to the larger N50 it generates, while ONT was able to characterize SVs smaller than 500 bp and detected 26,413of them.

Our analysis revealed differences in the types of structural variants (SVs) detected by the ONT pipeline and the OGM method (**Figure 3.A**). While the ONT pipeline, using Sniffles2 ^46^, only identified deletions (1,364) and insertions (2,501), OGM detected additional SV types, translocations (4) and inversions (49), alongside deletions (1,220) and insertions (2,924). OGM duplications don’t have defined confidence score, so they were excluded from the analysis. This observation suggests potential limitations of Sniffles2 for certain SV types, particularly translocations and inversions. A recent study by Bolognini and Magi ^47^ evaluating various SV callers within the ONT framework, suggests that alternative SV callers like SVIM^26^ or CuteSV ^27^ might outperform Sniffles2 in detecting such SV types, when used after the same aligner (minimap2). Based on these findings, we employed SVIM and CuteSV for SV detection in the tumor sample and compared the results of all three ONT callers to the OGM results (**Figure 3.A**). Notably, none of the tested ONT tools identified translocation breakpoints or inversions that met our quality filtering criteria (see methods).

**Figure 3.**
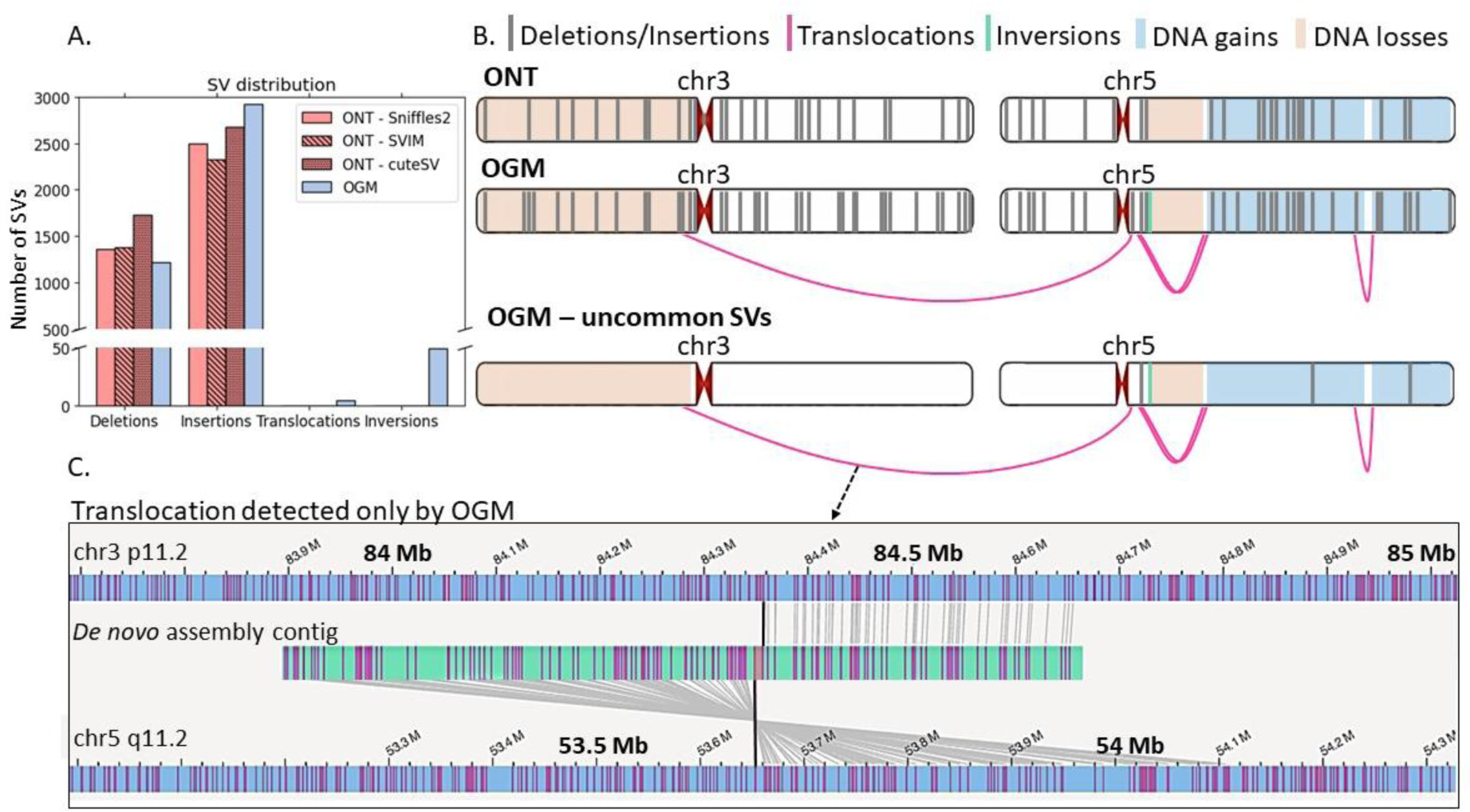
SVs detected by ONT and OGM. **A.** Comparison of number of SVs (≥500 bp) by type detected by OGM and three SV callers for ONT – Sniffles2, SVIM, CuteSV. **B.** Illustration SVs larger than 5 kbp on chromosomes 3 and 5, as detected by ONT (top panel) and OGM (middle panel). Bottom panel shows OGM SVs that do not appear in BNG’s dataset of healthy controls, hence potentially pathogenic. **C.** Interchromosomal translocation detected only by OGM. The light blue strips at the top and bottom represent the reference chromosomes 3 (top) and 5 (bottom), and the middle strip is a *de novo* assembled contig, composed of fragments mapped to chromosome 3 and inverted chromosome 5. Pink lines indicate CTTAAG barcode labels in the contig and reference. Black lines indicate translocation breakpoints.

**Figure 3.B** depicts chromosomes 3 and 5, two chromosomes that are frequently disrupted in ccRCC, with marks indicating the relative positions of SVs and CNVs larger than 5 kbp identified by ONT and OGM. SVs overlapping BNG’s list of N-base gaps in the reference or putative false positive translocation breakpoints (for *de novo* assembly, Solve 3.6.1) were masked for both methods. Out of the OGM-detected SVs, uncommon SVs not present in BNG’s database of healthy controls are separately plotted on the bottom. As seen also in **Figure 2**, the two methods detected DNA gain/loss events in these chromosomes and exhibited a high degree of concordance for insertions and deletions. OGM detected 3 possible inversions (similar locus), 3 intrachromosomal and 1 interchromosomal (**Figure 3.C**) translocation events. Notably, only 12 SVs identified by OGM in these chromosomes did not appear in BNG’s database of mapped healthy controls, indicating possible pathogenic SVs. Two of them were also found by ONT, and 10 of them are potential somatic variants not found in the normal sample adjacent to the tumor (of these, none were found by ONT). ONT SVs found in all chromosomes in the ccRCC tumor and the normal adjacent tissue are shown in figures **S2** (raw, unfiltered karyogram) and **S3** (processed circos plots). OGM SVs found in all chromosomes in the ccRCC tumor and the normal adjacent tissue are shown in figures **S4** (processed circos plots of all SVs), **S5** (processed circos plots of SVs not present in BNG’s database of healthy controls), and **S6** (BNG default display summarizing all results. SVs plotted are not present in BNG’s database of healthy controls).

### Epigenetic Analyses Methylation profiling

Modification calling is an attractive feature of ONT, that sequences native, unamplified DNA. Accordingly, modifications can be called directly from nanopore signal data, without needing chemical conversions (like in bisulfite sequencing, for example) ^48^. The current recommended basecaller for ONT, Dorado (https://github.com/nanoporetech/dorado), can call modified bases, and has ready-to-use models to call 5-methyl cytosine (5mC) and 5-hydroxymethyl cytosine (5hmC), on top of the four canonical bases.

Commercially-supported OGM only provides tools for analyzing genomic structure for cytogenetic applications. However, more layers of information may be multiplexed with the use of colors. Some of the BNG Saphyr systems contain three laser colors, two of them are for generating the genetic barcode and DNA backbone, and the third can be used with orthogonal chemistries to tag genomic features of choice, including epigenetic marks.

Unmethylated CpG sites, complementary to methylated (5mC) sites, can be specifically labeled by methyltransferase enzymes. We have recently applied an engineered CpG methyltransferase to address all unmethylated CpGs ^41,42^. However, the method was not yet validated for human methylome profiling and thus we used the previously validated reduced representation optical methylation mapping (ROM) ^14,16,21^. This method uses the methyltransferase M.TaqI, which directly transfers a fluorescent tag from a synthetic cofactor to an adenine base in the enzyme’s recognition sequence TCGA. However, if the CpG nested in this sequence motif is methylated or modified, the labeling reaction is blocked (**Figure 4.A**). Consequently, the DNA is labeled in all unmodified CpGs that are within TCGA sites. This reduced representation of the human methylome encompasses only ∼ 6% of the total CpGs but coincidently captures the majority of regulatory sites in the genome and has been shown to present a cell-type specific pattern ^14,16^.

**Figure 4.**
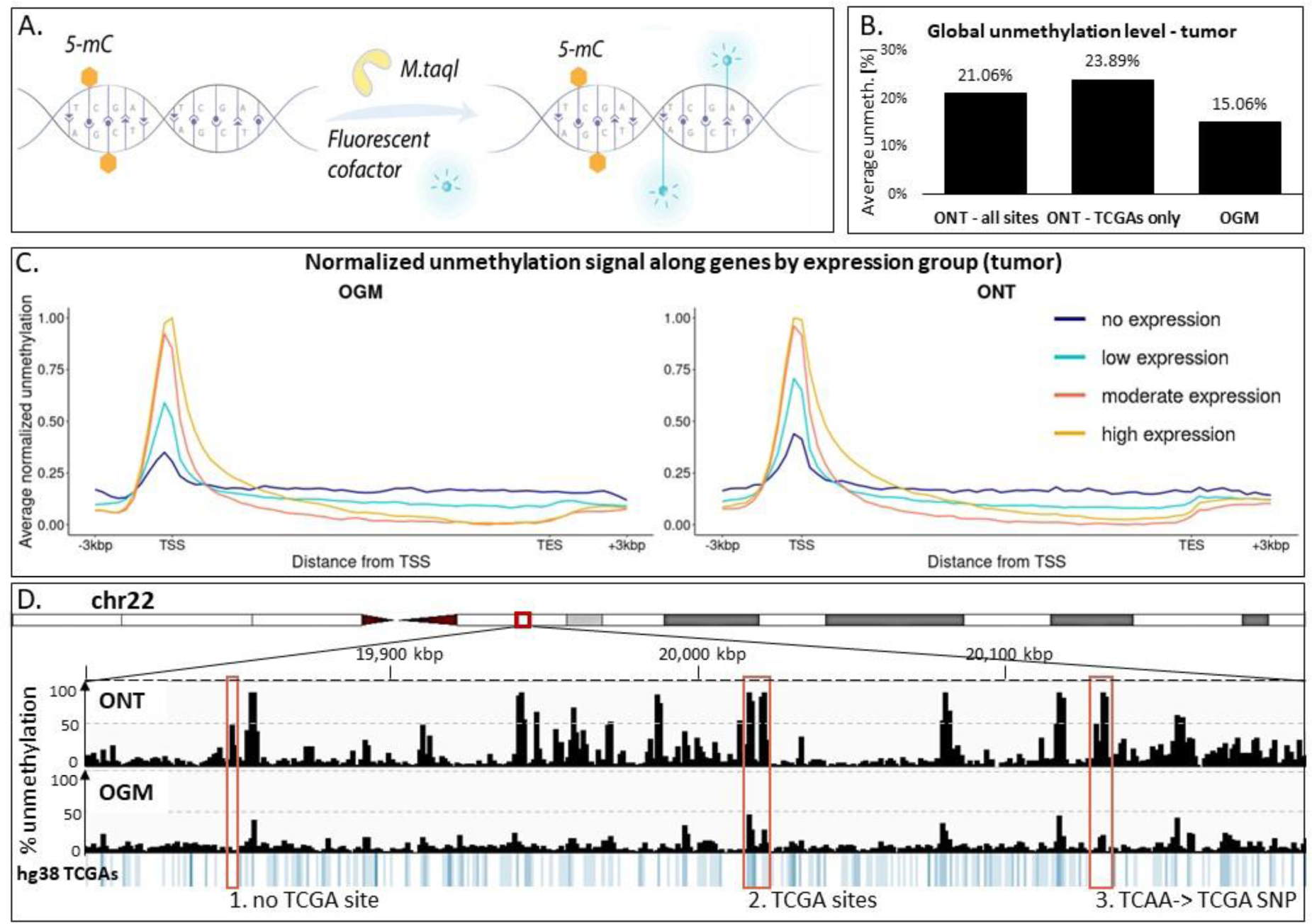
Unmethylation analysis. **A.** Fluorescent labeling scheme for unmodified CpGs embedded in TCGA motif for OGM. **B.** Average global unmethylation levels of a ccRCC tumor, as detected by ONT in all CpG sites, by ONT when restricted to TCGA-embedded CpG sites only, and by OGM (inherently marking only TCGA sites). **C.** ONT and OGM unmethylation signal, each normalized between 0 and 1, of the ccRCC tumor along aggregated genes. Genes were grouped based on their expression in ccRCC tumors. **D.** Unmethylation profiles of the ccRCC tumor by ONT and OGM along a region in chromosome 22, and the corresponding density of TCGA motif in the hg38 reference. Three red boxes mark three regions that differ in TCGA content (all regions contain CpG sites): 1: no TCGAs, there is a peak in ONT signal and not in OGM. 2: TCGAs are present, peaks in both methods. 3: no TCGAs in the reference, peaks in both methods due to a SNP.

To facilitate a direct comparison of methylation signals between the two methods, we transformed the ONT methylation values into unmethylation signals by presenting the complement to 1 of the calculated methylation level. We applied a minimum coverage threshold, requiring at least 5 reads for ONT and 20 reads for OGM. In order to account for the lower resolution of OGM, we calculated the average unmethylation signals in non-overlapping 1 kbp genomic windows. We compared the entire ONT methylome (all CpG sites) as well as a reduced ONT methylome (TCGA motif) to the reduced representation OGM signals (**Figure. 4.B**). Our analysis revealed a higher unmethylation signal in ONT compared to OGM. Interestingly, the difference persisted, and even slightly increased when we specifically analyzed TCGA-embedded CpG sites in ONT data. This suggests a potential underestimation of unmethylation by OGM, likely attributable to its lower optical resolution, rather than to the reduced representation approach. Consequently, multiple closely spaced TCGA sites might be erroneously merged into a single unit by OGM, leading to an underestimation of the overall unmethylation signal. Plots showing distances between adjacent TCGA sites, the number of TCGA sites in 1 kbp windows, and this number vs. the number of CpG sites in the same bins, are presented in **Figure S7**. **Figure 4.C** shows that despite absolute intensity differences, similar trends are seen in the normalized unmethylation profile generated along genes when grouped by their gene expression score in ccRCC tumors^29^. Both methods display the higher unmethylation signal around the transcription start site (TSS), which increases with gene expression. In contrast, the level in gene bodies is much lower and more similar among all expression levels. Un-normalized OGM and ONT levels of the tumor and normal adjacent tissue along genes are presented in **Figure S8**. The resolution and methylation representation differences become more apparent when zooming in to smaller regions of the genome. **Fig 4.D** shows the unmethylation profile of a ∼400 kbp region in chromosome 22q11.21. Three representative examples for methylation comparison are marked in red boxes that contain variable TCGA content (shown in blue in the lower panel). The leftmost box showcases a region lacking any TCGA sites in the reference genome. Consequently, the ONT plot exhibits a high unmethylation signal (indicating unmethylated CpGs), while the OGM profile shows no signal. The middle box highlights two adjacent bins with relatively high TCGA density, resulting in signal peaks by both methods. The rightmost box depicts a region lacking a reference TCGA site, yet the OGM profile displays a peak. Intriguingly, investigation of the corresponding ONT sequence revealed an A-to-G single nucleotide polymorphism (SNP), creating a new TCGA site recognizable by the M.TaqI enzyme, thus explaining the observed OGM signal.

### hydroxymethylation profiling

5-hydroxymethyl cytosine (5hmC), the first oxidation product of 5-methylcytosine, is another important modification that was linked to gene regulation, development and disease, predominantly cancer ^49,50^. 5hmC calling has recently been integrated to the ONT basecaller Dorado (https://github.com/nanoporetech/dorado). Identification of modifications in ONT data relies on machine learning techniques. This process involves training and validating models using reference data encompassing the modification across diverse sequence contexts. Such reference data can be obtained by identifying the modification using established methods or in-situ approaches ^17^. However, obtaining high-quality, genome-wide reference data specifically for 5hmC modifications remains a significant challenge due to its cost and complexity. This, in turn, limits the ability to comprehensively train and assess the performance of 5hmC callers for ONT data, and it hasn’t been benchmarked and peer-reviewed to date.

Optical mapping of 5hmC was introduced several years ago based on the fluorescent labeling of 5hmC residues ^15,21^. 5hmC is directly labeled in a process that involves the enzymatic attachment of an azide-modified glucose moiety from a synthetic cofactor ^51,52^ (UDP-6-N3-Glu) to the hydroxyl group of 5-hmC, followed by a click reaction that connects a fluorophore-bound alkyne to the azide-labeled 5-hmC ^53,54^ (**Fig 5.A**). **Fig. 5.B** shows the average genome-wide 5hmC signal in the ccRCC tumor sample and the normal adjacent tissue, as was detected by both methods. Consistent with published reports indicating a global reduction of 5hmC in various cancers ^50,55,56^, both methods revealed a ∼3-fold decrease in 5hmC levels in the tumor compared to the adjacent normal tissue. This time, OGM detected higher absolute levels of 5hmC compared to ONT. As the labeling scheme used to tag 5hmC residues in the OGM experiment has no false positives, and was validated with LC-MS/MS in previous work ^15^, we hypothesize that there is an underestimation of 5hmC calls by the ONT model due to incomplete training sets and challenging sequence contexts. Albeit showing different absolute levels of 5hmC, the modulation of 5hmC level along gene bodies, as well as the increase in signal as a function of gene expression, can be seen by both methods (**Figure 5.C** and **Figure S9**.). The 5hmC profile generated by both methods and displayed in **Figure 5.D** reveals a broadly correlated profile, but with distinct amplitude variations between the different datasets, in line with the average global levels. **Figure 5.E** shows an example of a large repetitive element containing a group of genes from the *GAGE* family, poorly represented in the hg38 reference (the entire array spans ∼190 kbp in the reference, with a gap within these coordinates) ^57^. Long molecules spanning the entire uncharacterized region in OGM aided in assembling a contig of the full repetitive element, and the 5hmC tags on these molecules provided the 5hmC profile along the unknown region. The panel also depicts a 5hmC-containing single molecule (digitized) and the average 5hmC signal along the contig. Epigenetic characterization of this region by ONT was not possible due to the shorter molecules that could only penetrate several thousand bases into the ENCODE blacklist-masked *GAGE12* region ^58^.

**Figure 5.**
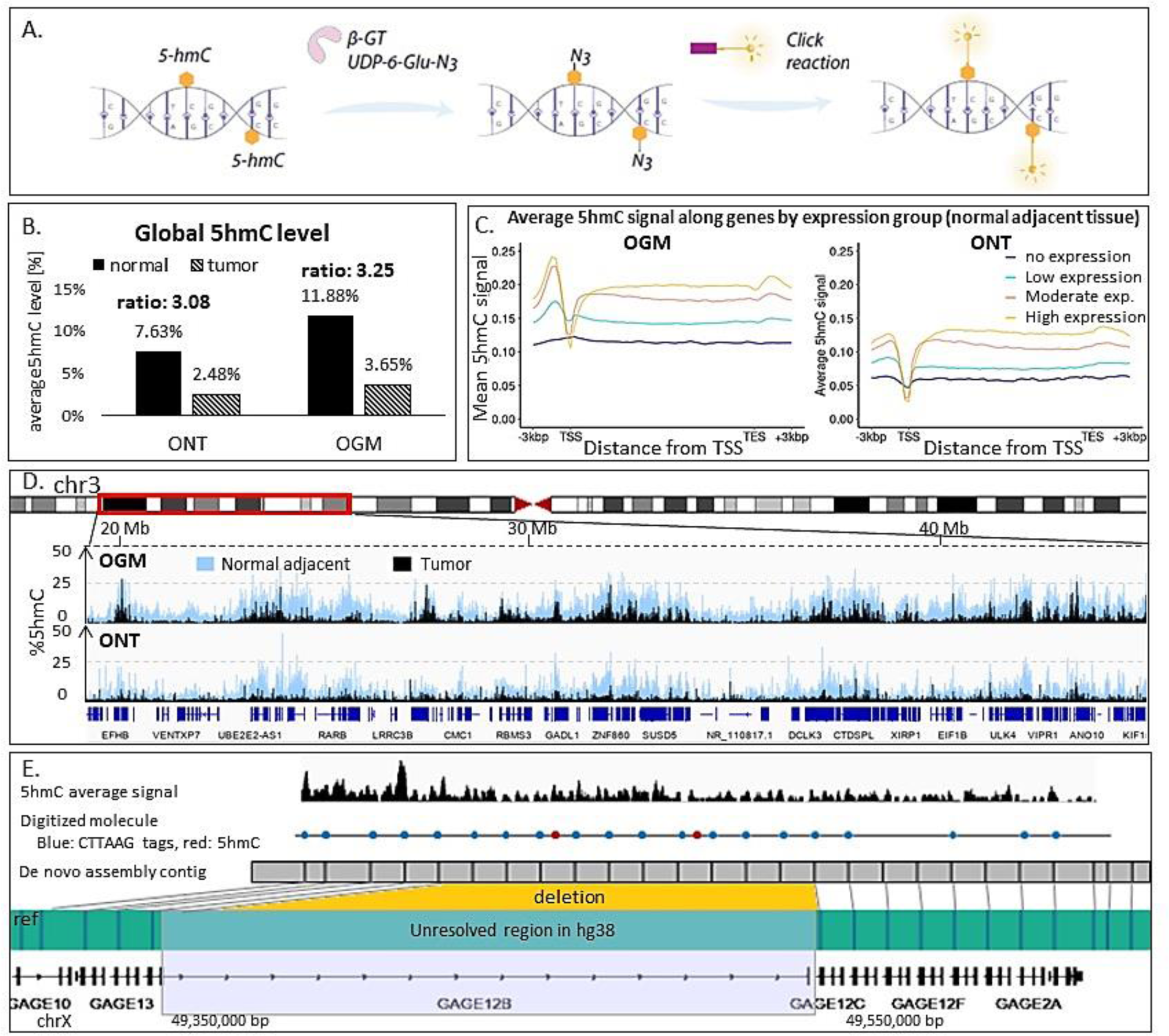
5hmC analysis. **A.** Direct fluorescent labeling of 5hmC for OGM. **B.** Global 5hmC levels of a ccRCC tumor and a normal adjacent tissue, as detected in OGM and ONT. **C.** OGM and ONT 5hmC signal of a normal kidney tissue adjacent to a ccRCC tumor along aggregated genes. Genes are grouped based on their expression in kidney tissues adjacent to ccRCC tumors. **D.** 5hmC profiles of a ccRCC tumor and a normal adjacent tissue generated by OGM and ONT along a ∼25 Mb region in chromosome 3. **E.** A repetitive sequence element in chromosome X, poorly characterized in the hg38 reference (green strip; Blue lines on it indicate the genetic barcode labels) above it, a *de novo* assembled OGM contig (gray) spanning the entire repeat array, indicating a deletion compared to hg38. The region spans genes from the *GAGE* family, and the gapped region contains the gene *GAGE12B*. Above it, a digitized single OGM molecule, with genetic barcode labels (blue) and 5hmC labels (red). Above it, is the average 5hmC signal along the contig.

## Discussion

BNG and ONT now offer tools that aim to unveil the complexity of aberrated genomes and replace many cytogenetic workflows. Both companies have developed dedicated toolkits for variant calling. To navigate this evolving landscape, this report offers an objective comparison of the two methods, delving into the data types accessible with each technology, and the capabilities of their respective analytical tools, recognizing these tools as crucial for generating reports with clear clinical relevance. In this respect, BNG is more clinically oriented in the cytogenetic space, with pipelines and reports that are aligned with clinical needs. BNG additionally compiled a substantial reference database of healthy controls. This enables the filtering of non-pathogenic findings.

At the karyotype level, the methods conform, and both are capable of providing reliable copy number evaluations. Nevertheless, slight differences in copy number can be observed (**Figures 2.A** and **S1**) and are attributed to the higher resolution of ONT. As for detecting structural variations in variable sizes, a trade-off between resolution and read-length was observed: both methods called a high percentage of mid-range SVs (1-10 Kbp). ONT, owing to its single-base resolution, performed better in detecting small SVs. OGM on the other hand, showed a clear advantage in detecting large SVs (larger than 10 kbp) with over 70% of large SVs (including translocations) not detected by ONT. Despite the low information content of OGM, the high coverage and N50 of these measurements increase the chance of detecting challenging structural aberrations. ONT’s resolution advantage is also significant for reporting the SVs’ breakpoints at a single bp resolution. This gives access to single nucleotide polymorphisms and short insertions and deletions that OGM is blind to. On the other hand, large or complex SVs are challenging for ONT under the experimental N50 and coverage. This holds true also for rarer variants that are detected by OGM due to the high relative genome coverage generated. Translocations and inversions, were not reported by neither of the ONT SV callers Sniffles2, SVIM and CuteSV, while OGM called inversions, three intrachromosomal translocations on chromosome 5 and one interchromosomal translocation between chromosomes 3 and 5. Per dollar, the genomic coverage generated by OGM is higher than that of ONT, opening a window to detect low-frequency variants and more resilience to sample heterogeneity. However, in this experiment, we did not meet the recommended coverage for running BNG’s “rare variant pipeline*”* (300x is recommended for high sensitivity to low frequency variants ^59^), therefore we performed the pipelines of “*De novo* assembly” and “variant annotation pipeline*”*, instead. The choice of analytical tools significantly influences the insights extracted from data generated by both methods. While we employed recommended tools optimized for our data type and coverage, we note that these tools have inherent limitations that potentially extend beyond purely technological constraints ^47,60,61^.

As for epigenetics, ONT can now call methylated CpGs from native DNA, together with generation of genetic data, an obvious advantage compared to OGM. OGM users that seek methylation information have to fluorescently tag the epigenetic modifications prior to data acquisition. These additional labeling steps are not commercialized by BNG and are not supported by the company. Methylation mapping extent is confined by the ability of the methyltransferase enzyme selected for this procedure and the density of its recognition sites. The enzyme M.TaqI, described here, efficiently labels CpG sites nested within the TCGA motif ^14^. This provides a reduced representation of the unmethylome. These recognition sites make up ∼6% of the CpG sites in the human reference, with correlating methylation states in many important regions of the genome ^14^, but inherent reduced representation limitations apply, in addition to constrains added by the difficulty to resolve adjacent labels due to optical resolution (diffraction limit). Additionally, the indirect labeling done by methyltransferase enzymes, pointing unmodified sites, can’t distinguish methylation from other cytosine modifications and is subjected to labeling efficiency, thus is inferior to direct methylation calling. Our analysis showed that global trends, such as correlation of signal with gene expression group, persisted, while locus specific signals depend on TCGA representation. This comparison highlights OGM’s limitations in methylation calling compared to ONT. However, to date, the picture for 5hmC presents a different scenario. In this case, the fluorescent labeling added to OGM, while also external and not supported by BNG, directly labels 5hmC residues and not complementary sites ^15^. Similarly to methylation calling, ONT enables 5hmC identification together with canonical basecalling without additional experimental steps, but some differences have to be considered. As the process of modification calling relies on machine learning, model training is a crucial step for accurate identification of 5hmC. This step requires comprehensive reference data covering the modification in all possible sequence contexts and distinguishing it from other cytosine modifications to assure accurate calls. Unfortunately, unlike for methylation, obtaining high-quality genome-wide reference data for 5hmC is still challenging and expensive, and might limit the comprehensiveness of the training data, thus affecting the performance of 5hmC calling models. This might explain the lower 5hmC levels called by ONT compared to OGM, seen in our comparison, and suggest that the ONT model currently underestimates the density of 5hmC and misses many of the modified bases.

To conclude, selecting the most suitable platform hinges on a clear understanding of the data requirements dictated by the clinical or research question. To this end, a thorough understanding of the data generated by each platform, alongside the strengths and limitations of their respective analytical toolkits is needed.

## Funding

This work was supported by the European Research Council consolidator [grant number 817811]; Israel Science Foundation [grant number 771/21]; The National Institute of health/The National Human Genome Research Institute (NIH/NHGRI) [grant number R01HG009190].

The authors declare no competing interests.

## Author contribution

**SM**: Methodology, Software, Formal analysis, Investigation, Writing - Original Draft; **ZT**: Investigation, Formal analysis; **TDZ, YM, JD**, **GN**: Investigation; **RH**: Writing - Review & Editing; **YG**: Resources; **DO**: Resources; **BD**: Resources; **HBF**: Writing - Review & Editing; **AG**: Conceptualization, Methodology, Software, Formal analysis, Investigation, Writing - Original Draft; **YE**: Conceptualization, Writing - Original Draft.

## Data availability

Data generated during this study will be made publicly available upon acceptance for publication.

## Declaration of generative AI and AI-assisted technologies in the writing process

During the preparation of this work the authors used Gemini in order to improve language and readability. After using this tool, the authors reviewed and edited the content as needed and take full responsibility for the content of the publication.

## Notes

### Competing Interest Statement

The authors have declared no competing interest.

### Summary of Updates

Revised manuscript. Main changes in figures 2,3,4. Supplementary information added.

