## Supplementary material for "Long-Read Structural and Epigenetic Profiling of a Kidney Tumor-Matched Sample with Nanopore Sequencing and Optical Genome Mapping": S1, S2, S3, S4, S5, S6, S7, S8, S9

### Supplementary Information

#### Supplemental Figures

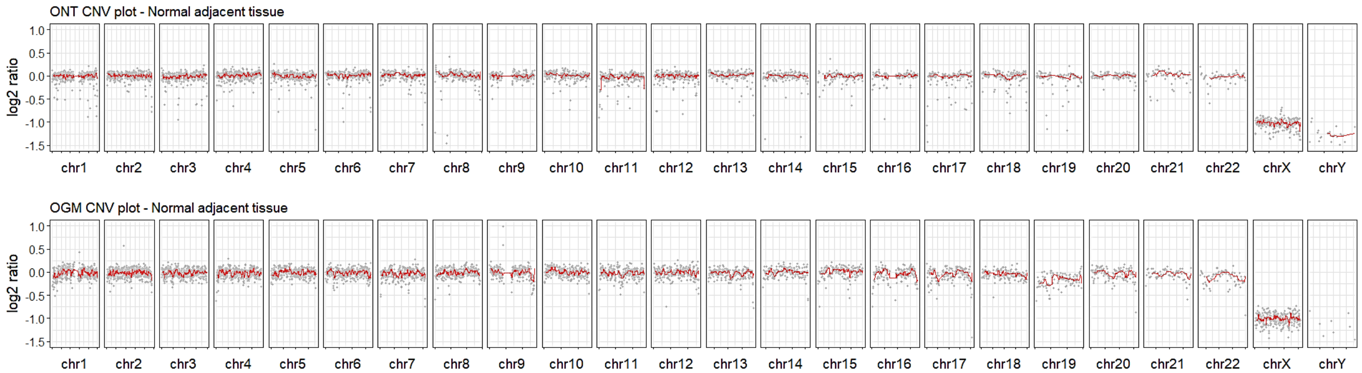


**Figure S1.** CNV plots ($\log_2$ of the copy ratio) generated from ONT (top) and OGM (bottom) data of a normal kidney tissue adjacent to a ccRCC tumor.

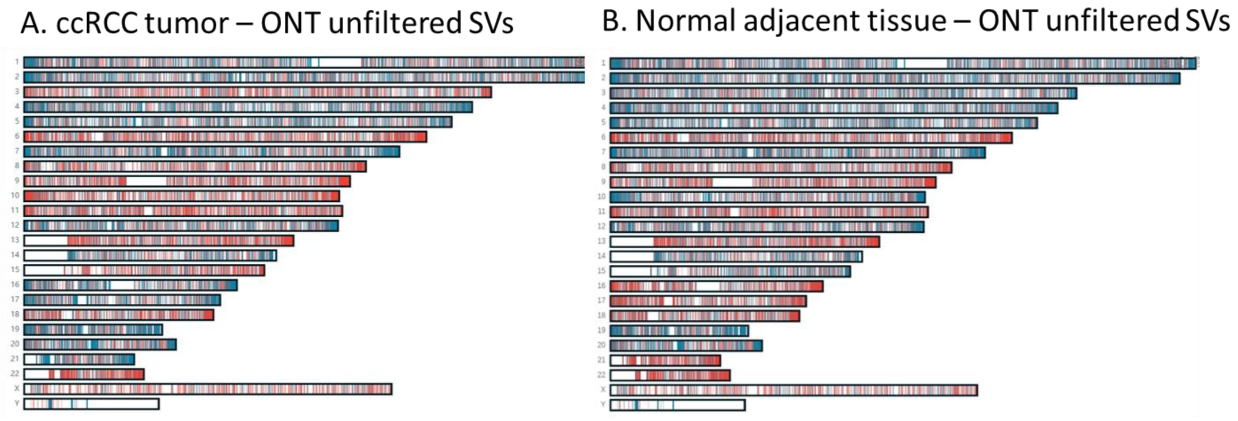


**Figure S2.** Chromosomal hotspots of structural variation called by Sniffles2 from ONT data. Karyograms contain raw, unfiltered SVs as plotted by the EPI2ME report. Deletions are shown in blue and insertions in red (Other SV types weren't called). **A.** SVs in the ccRCC tumor sample. **B.** SVs in the normal tissue adjacent to the ccRCC tumor.

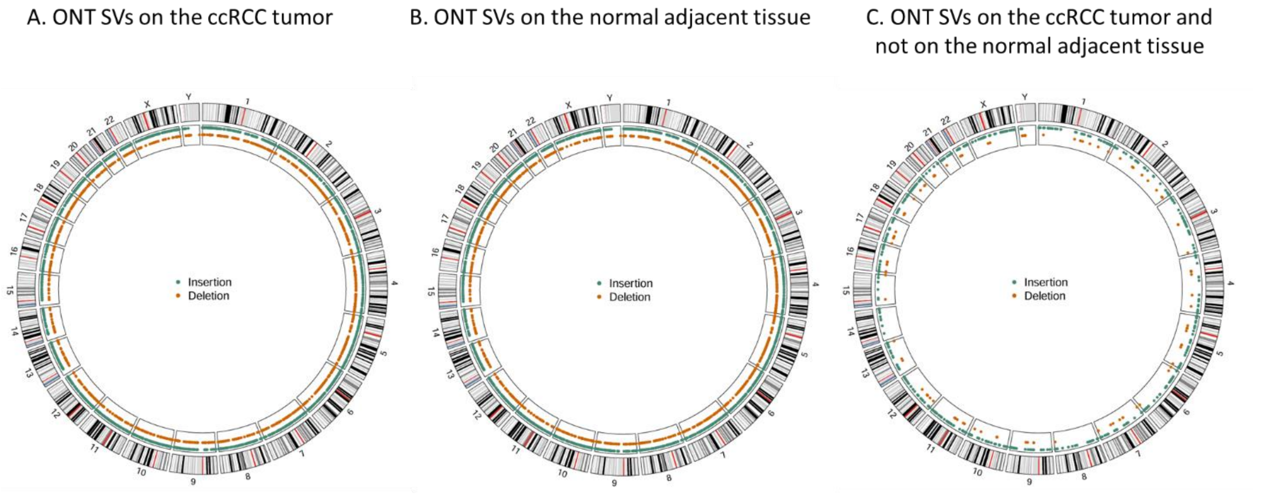


**Figure S3.** Circos plots showing chromosomal hotspots of structural variations larger than 500 bp called by Sniffles2 from ONT data. SVs displayed are supported by at least 5 reads, were labeled with ‘pass’ filter and do not overlap BNG’s list of N-base gaps in the reference or putative false positive translocation breakpoints (for “*de novo* assembly”, Solve 3.6.1). Deletions are shown in orange and insertions in green (Other SV types weren’t called). Plots were created with the R package Circlize [1]. **A.** SVs on the ccRCC tumor. **B.** SVs on the normal adjacent tissue. **C.** SVs on the tumor tissue not overlapping SVs on the normal adjacent tissue.

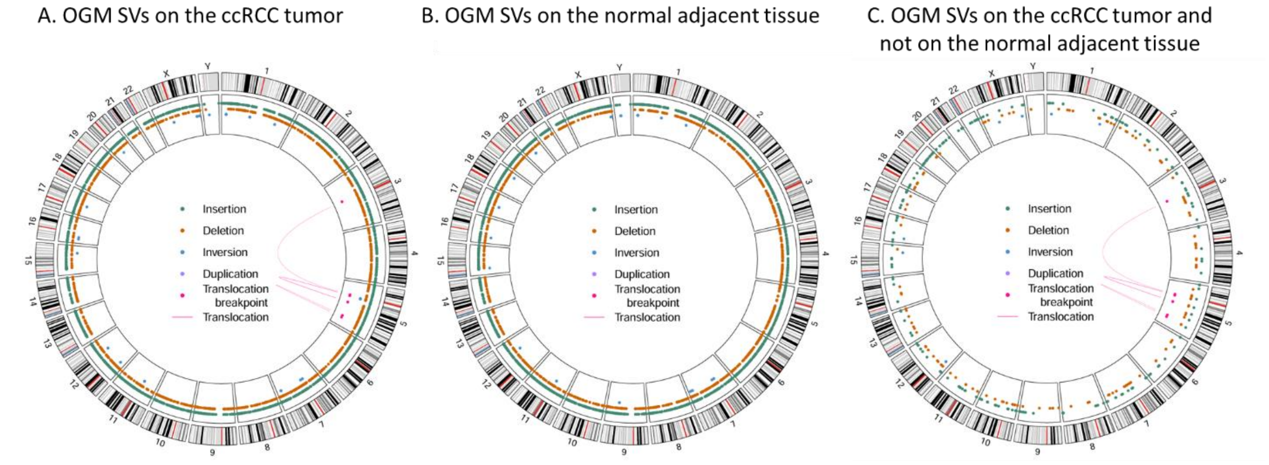


**Figure S4.** Circos plots showing chromosomal hotspots of structural variations called by OGM. SVs displayed have confidence score ≥ 0.5 and do not overlap BNG’s list of N-base gaps in the reference or putative false positive translocation breakpoints (for “*de novo* assembly”, Solve 3.6.1). Insertions are shown as green dots, deletions in as orange dots, inversions as blue dots, duplications (not present due to confidence filtration) as purple dots, translocation breakpoints as pink dots, and translocations as pink lines connecting the breakpoints. Plots were created with the R package Circlize [1]. **A.** SVs on the ccRCC tumor. **B.** SVs on the normal adjacent tissue. **C.** SVs on the tumor tissue not overlapping SVs on the normal adjacent tissue.

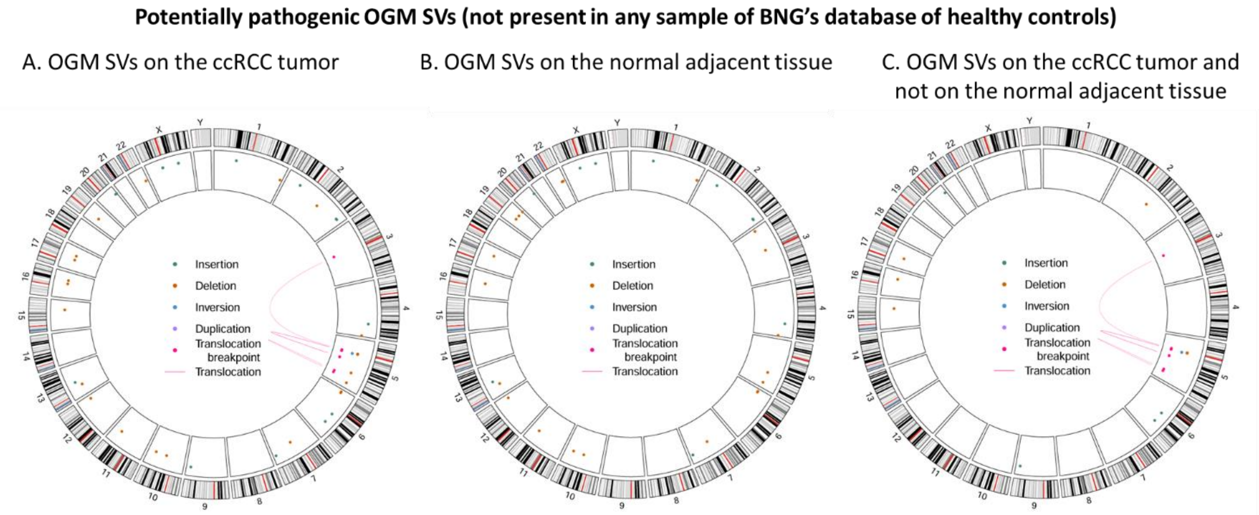


**Figure S5.** Circos plots showing chromosomal hotspots of structural variations called by OGM, and not present in BNG's database of healthy controls. SVs displayed have confidence score ≥ 0.5 and do not overlap BNG's list of N-base gaps in the reference or putative false positive translocation breakpoints (for "*de novo* assembly", Solve 3.6.1). Insertions are shown as green dots, deletions in as orange dots, inversions as blue dots, duplications (not present due to confidence filtration) as purple dots, translocation breakpoints as pink dots, and translocations as pink lines connecting the breakpoints. Plots were created with the R package Circlize [1]. **A.** SVs on the ccRCC tumor. **B.** SVs on the normal adjacent tissue. **C.** SVs on the tumor tissue not overlapping SVs on the normal adjacent tissue.

**Potentially pathogenic OGM SVs and CNVs – BNG output**

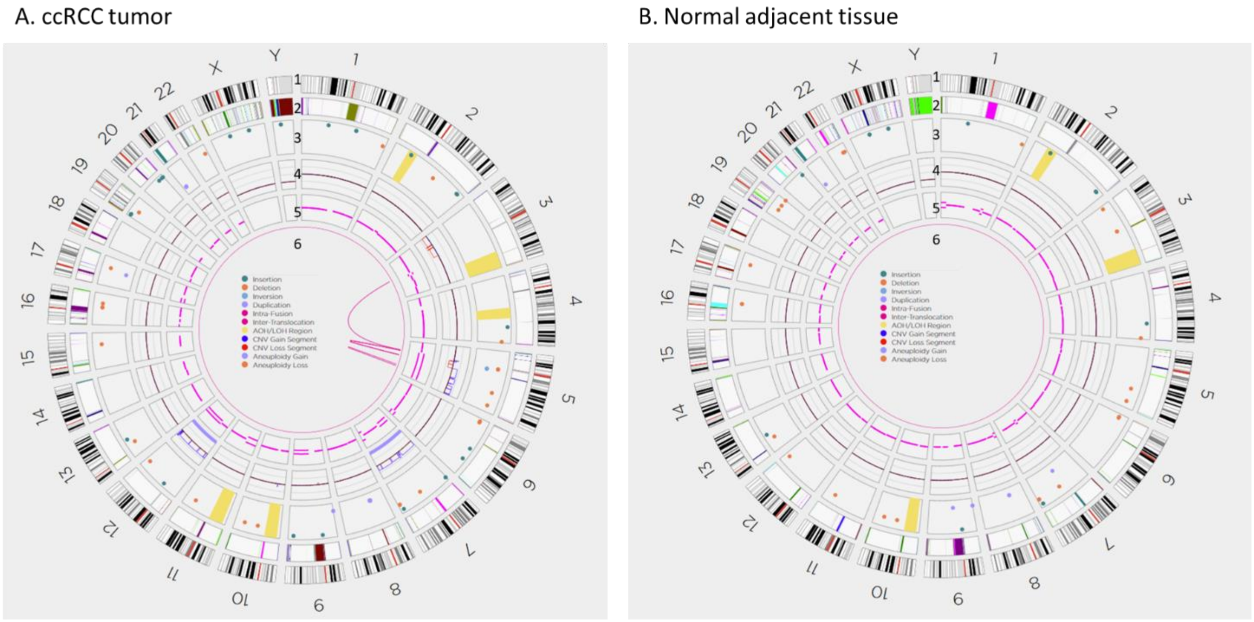


**Figure S6.** Circos plots showing results of OGM *de novo* assembly. The plots depict 6 information layers. From outmost to inner: 1: cytobands, 2: masked locations in CNV analysis, 3: SVs not present in BNG's database of healthy controls, appear as dots according to the following color code: green: insertions; orange: deletions; light blue: inversions; purple: duplications. Yellow bands indicate absence/loss of heterozygosity. SVs confidence filter is according to BNG's defaults: confidence ≥ 0 for insertions and deletions, ≥ 0.7 for inversions. Confidence score for duplications is undefined. 4: Copy number baseline and copy number variations. DNA gains are in blue, losses in red, aneuploidy gain in purple and aneuploidy loss in orange. 5: Variant allele frequency. 6: Translocations, including intra chromosomal fusions (confidence ≥ 0.3) and for inter chromosomal translocations (confidence score ≥ 0.65). Translocations shown are not present in BNG's database of healthy controls. **A.** ccRCC tumor. **B.** normal adjacent tissue.

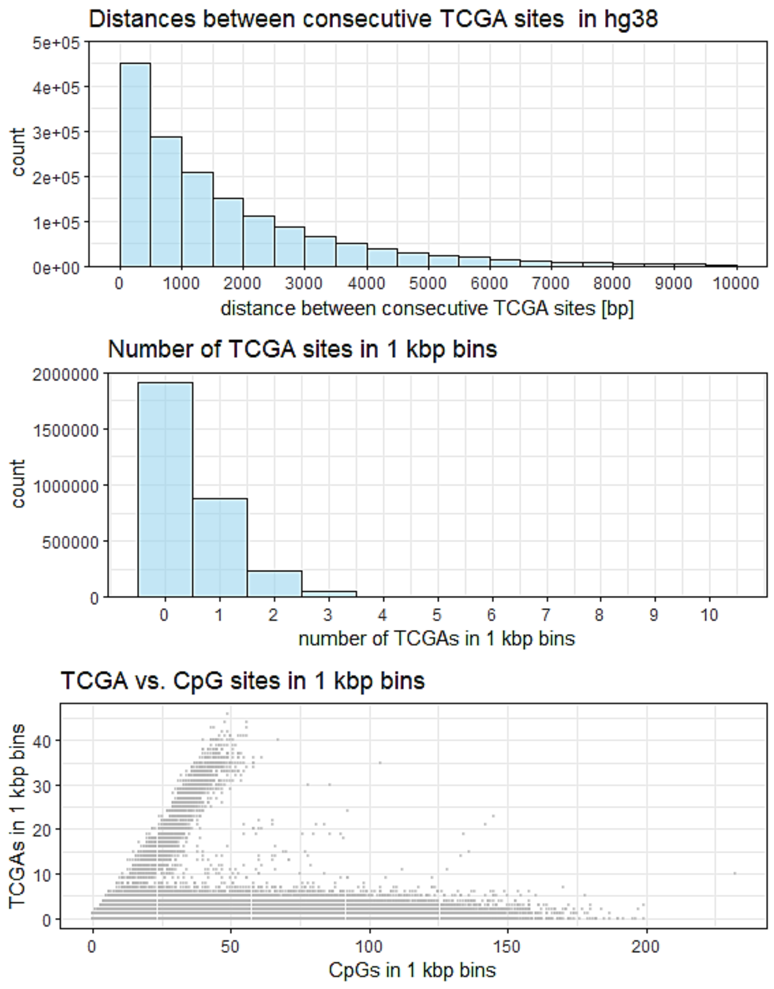


**Figure S7.** TCGA sequence motive. **A.** Distance between consecutive TCGA sites. 25% of sites are located within 500 bp of the previous site. 50% of sites are located within 1500 bp of the previous site. 75% of sites are located within 2500 bp of the previous site. **B.** Number of TCGA sites in 1000 bp genomic windows. **C.** TCGA vs. CpG content in 1 kb genomic windows.

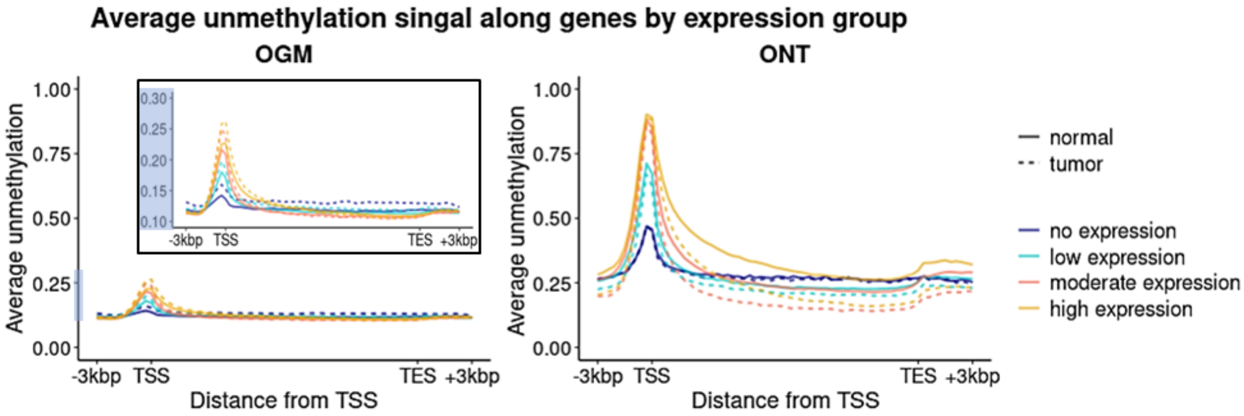


**Figure S8.** Unnormalized OGM (left) and ONT (right) unmethylation signal of the ccRCC tumor and normal adjacent tissue along genes aggregated based on their expression in corresponding tissues (ccRCC tumors/ normal tissues adjacent to ccRCC tumors).

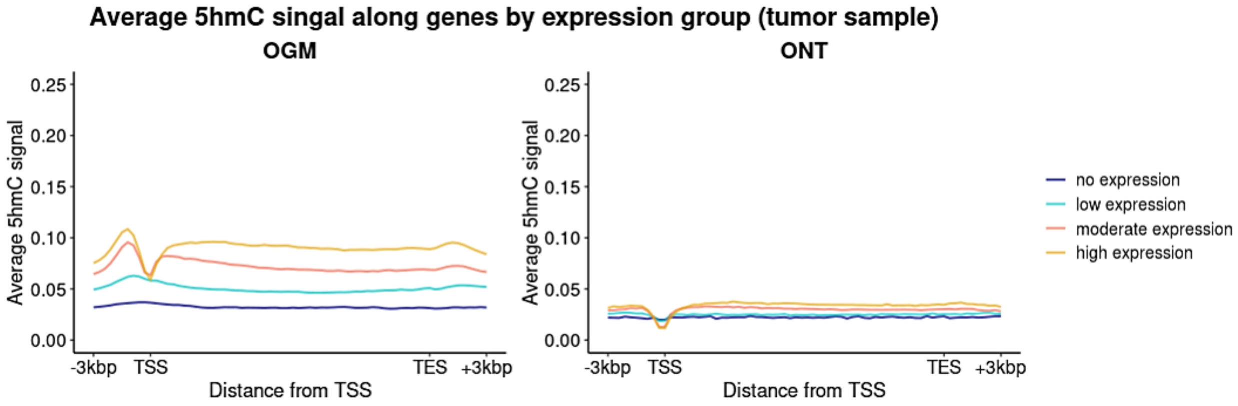


**Figure S9.** OGM (left) and ONT (right) 5hmC signal of a ccRCC tumor along aggregated genes. Genes are grouped based on their expression in ccRCC tumors.
